# *Culicoides* allergens expressed in insect cells induce sulfidoleukotriene release in peripheral blood leukocytes from horses affected with insect bite hypersensitivity

**DOI:** 10.1101/2025.01.27.635044

**Authors:** S Jonsdottir, SB Stefansdottir, J Mirkovitch, A Ziegler, S Torsteinsdottir, E Marti

## Abstract

**Introduction:** Insect bite hypersensitivity (IBH) is an IgE mediated dermatitis in horses caused by bites of *Culicoides* spp. The allergens are salivary gland proteins from these insects and nine major allergens from *Culicoides obsoletus* have been identified and expressed in *E. coli*. However, proteins expressed in procaryotic systems have limitations in cellular assays, particularly in functional assays assessing the allergen-induced release of mediators *in vitro*, such as sulfidoleukotrienes (sLT) from basophils. Aims of the study were to produce functional *Culicoides* allergens in insect cells, to assess their allergenicity using a sLT release assay and to relate the sLT release with IgE sensitization to the respective allergens using ELISA.

**Methods:** Eight major *Culicoides obsoletus* allergens (Cul o 1P, Cul o 2P, Cul o 3, Cul o 5, Cul o 7, Cul o 8, Cul o 9 and Cul o 11) were expressed in insect cells and purified. sLT release from peripheral blood leukocytes (PBL) following stimulation with the eight *Culicoides* allergens was measured in 28 IBH-affected and 24 healthy control horses. Allergen-specific serum IgE levels was determined by ELISA.

**Results:** The eight major allergens were successfully expressed in insect cells and purified. All allergens induced a significantly higher sLT release from PBL of IBH-affected horses compared to healthy controls. There was a high correlation and substantial to excellent agreement between sLT release and serum IgE levels for six *Culicoides* allergens, while for two the agreement was moderate. Positivity rates in IBH horses were usually higher in IgE serology but more false positive results were obtained. The allergens performing best in both assays were Cul o 3, Cul o 8 and Cul o 9, with very high specificity and good sensitivity.

**Discussion:** Insect cell expressed *Culicoides* recombinant allergens are functionally relevant and will open new opportunities for the study of Culicoides hypersensitivity not only in horses, but also potentially in human patients or other species. They will also greatly improve IBH diagnostics using cellular assays as well as IgE serology.

## 1 Introduction

Allergy to bites of insects of the genus *Culicoides* is mostly known in horses (reviewed in 1, 2), but few reports indicate that it can also occur in humans (3, 4) and other species (5–7). Culicoides hypersensitivity, in horses also named insect bite hypersensitivity (IBH) or summer eczema is a type I, IgE-mediated allergy, and is the most common allergic disease in horses. It is caused by bites of insects of the genus *Culicoides* (midges). IBH is characterized by seasonal, severe itching, resulting in loss of hair, skin lesions, and sometimes secondary infections (8). All breeds of horses can be affected but the prevalence differs based on genetic and environmental factors. Horses of the Icelandic breed which are born in Iceland are especially prone to develop IBH after export to continental Europe. More than 50% of them develop IBH within a few years after export. This is due to lack of exposure to *Culicoides* bites at early age as *Culicoides* that feed on horses are not endemic in Iceland (9).

Diagnosis of IBH is mostly based on clinical history and typical clinical signs. Until recently, commercially available IgE serology was based on ELISA technology using *Culicoides* whole body extracts (WBE), derived from laboratory-bred *Culicoides nubeculosus*. This resulted in a low specificity and sensitivity of IgE serology (10). A sLT release assay (Cellular Antigen Stimulation Test (CAST); Bühlmann labs) with *Culicoides nubeculosus* WBE showed a much better specificity but a still moderate sensitivity of 80% for *in vitro* diagnosis of IBH (11). In a study by van der Meide et al. *Culicoides* whole body extract form the three different *Culicoides* species, *C. obsoletus* (CO), *C. nubeculosus* (CN) and *C. sonorensis* (CS) were compared in a histamine release assay and IgE serology (12). *C. obsoletus,* the most common species in the environment of the tested horses, showed the best performance in both tests.

However, the availability of WBE from *C. obsoletus* is limited as they cannot be bred in the laboratory and must therefore be caught in the wild, prohibiting its use at a larger scale. Furthermore, using WBE has limitations, as they are difficult to standardize for the presence of relevant allergens and the allergen content is low, as the causative allergens for IBH are salivary gland proteins. Thirty allergens have been identified from the three different *Culicoides* spp., *C. sonorensis* (13), *C. nubeculosus* (14), and *C. obsoletus* (15–17). They have all been expressed in *E. coli* and some in insect cells and barley (17–19). Nine of these allergens, (Cul o 1P, Cul o 2P, Cul o 3, Cul o 5, Cul o 7, Cul o 8, Cul o 9, Cul o 10 and Cul o 11) were shown to be major allergens for IBH using IgE serology with microarray technology. While their IgE-binding capability has been documented in various studies and using a larger number of sera (15–17, 20), the functional relevance has only been investigated for Cul o 5 and Cul o 7 by skin tests on a small number of horses (15). Functional *in vitro* assays such as basophil activation tests (BAT) or histamine or sLT release assays provide a functional assessment of the allergic response, unlike IgE serology that only measures sensitization.

These functional tests measure the ability of allergen-specific IgE to activate basophils, more closely replicating *in vivo* allergic reactions. These functional assays allow to potentially predict clinical reactivity and severity of allergic reaction and are useful to monitor the response to allergen immunotherapy (AIT) (21, 22). BAT is presently not available for the horse due to the lack of equine specific reagents. Histamine and sLT release assays are thus still used for this species, and a high correlation between these assays has been demonstrated previously (23).

Allergens produced in *E.coli* have been shown to work well in serology while showing low performance in cellular assays (18, 24). Unpublished work showed that *E.coli* expressed *Culicoides* allergens usually induce no, or only very low release of sLT from PBL of IBH-affected horses in CAST (unpublished data, Eliane Marti). Soldatova et al. showed that insect cell expressed bee allergen was functionally similar to the native protein while the same protein expressed in *E. coli* showed only one third of the enzymatic performance (25). While earlier studies showed that the enzymatic/catalytic activity of allergens such as recombinant PLA is not a requirement for allergenicity in the effector phase, incorrect folding of recombinant protein resulted in a total loss of allergenic potency (26). For some allergens post-translational modifications, which are missing from *E.coli* expressed proteins, are essential for the correct three-dimensional conformation of the molecule, its biologic activity and the correct conformation needed for its allergenicity (27). This highlights the importance of appropriate expression system based on the downstream application.

Our aim was to produce major *Culicoides* allergens in insect cells, evaluate their functional relevance for equine IBH using a Cellular Antigen Stimulation Test, and compare the resulting sLT release with IgE sensitization measured by ELISA.

## 2 Material and Methods

### 2.1 Cloning and expression of *Culicoides* allergens

Eight *Culicoides obsoletus* allergens (Table 1) were expressed in insect cells with the Bac-to Bac Baculovirus expression system (Thermo Fischer Scientific, Waltman, USA) according to manufacturer’s procedure. The genes encoding *Cul o 5* to *Cul o 9* and *Cul o 11* were codon optimized for expression in insect cells (GenScript, Rijswijk, NL) and were cloned into various expression vectors including pFastBac-1, pFastBac-HBM-TOPO and pI-secSUMOstar. Recombinant baculoviruses were harvested following a transfection into Sf-9 cells, cloned with limiting dilution, and amplified in Sf-9 cells at 27°C in closed culture in SF-900™II Serum Free Medium (Gibco® by Life Technologies™, Thermo Fischer Scientific), supplemented with 100 IU/mL penicillin, 100 µg/mL streptomycin, 1% fetal bovine serum (Gibco® by Life Technologies™, Thermo Fischer Scientific). Protein production was performed in High five insect cells; 200 mL of 3×10^6^ cells/mL (in the same medium as for Sf-9 cells without serum) were infected with 1-2 m.o.i. of third passage of cloned viruses and incubated at 15°C at 120 rpm in 500 mL Erlenmeyer flasks in an orbital shaker for 6-10 days, depending on the protein or until sufficient cytopathy. Cells were harvested by centrifugation at 515 x g for 12 min at RT, the pellets snap frozen in liquid nitrogen and stored at −80°C.

**Table 1:**
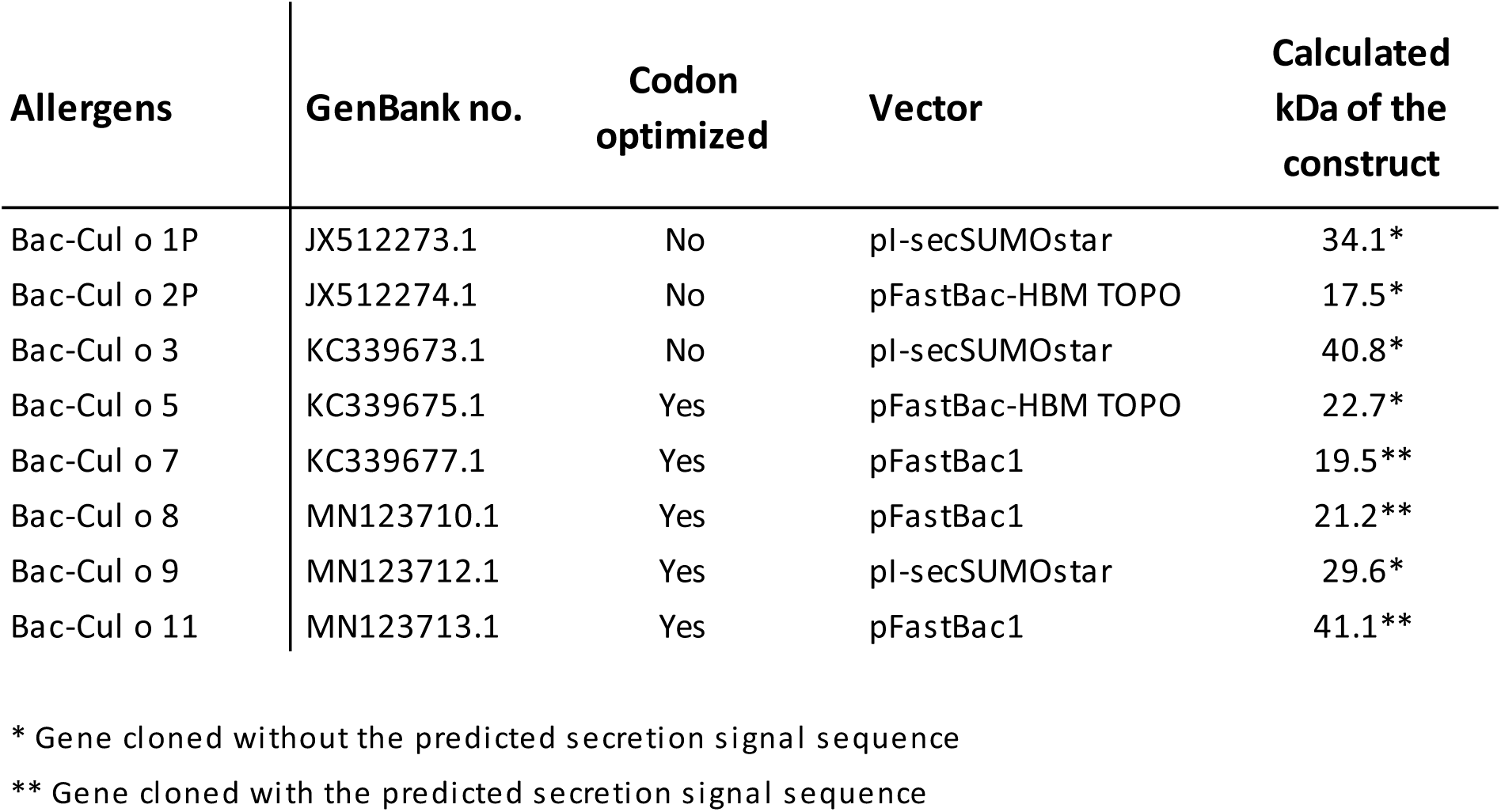
List of the eight major *Culicoides obsoletus* allergens. expressed in High five insect cells with the Bac-to Bac Baculovirus expression system and their calculated size in kDa based on the amino acid sequence of the construct.

### 2.2 Production of antibodies specific for *C. obsoletus* allergens

Polyclonal antibodies (pAb) specific for Cul o 1P, Cul o 2P, Cul o 5, Cul o 8, Cul o 9 and Cul o 11 were produced at the Institute for Experimental Pathology, University of Iceland, Keldur (Reykjavik, Iceland) or at ArcticLas (Reykjavik, Iceland) according to Schaffartzik et al. 2011 (1). *C. obsoletus* allergens (17) produced in *E.coli* and purified were used to immunize the mice, except for Cul o 8 where Bac-Cul o 8 was used.

Monoclonal antibody (mAb) specific for Cul n 1 was produced in mice with the hybridoma technique according to Köhler and Milstein 1975 (28) at the Institute for Experimental Pathology, University of Iceland, Keldur (Reykjavík, Iceland) in accordance with Janda et al. 2012 (29) and Jonsdottir et al. 2017 (19). *E.coli* produced and purified Cul n 1 was used to immunize mice (30).

The production of polyclonal and monoclonal antibodies was approved by the National Animal Research Committee of Iceland.

### 2.3 Purification of *Culicoides* allergens

All but one (Cul o 1P) of the eight His-tagged *Culicoides obsoletus* proteins were purified under native conditions using HIS-select® HF Nickel Affinity gel (Sigma-Aldrich, Merck, Burlington, USA) according to manufacturer’s procedures. In short: the 200 mL High five culture was split in 4 and a pellet from 150×10^6^ cells used for purification at each time. The pellet was resuspended in 8 mL lysis buffer (50 mM NaH_2_PO_4_xH_2_O, 150 mM NaCl, 1% IgePal, pH 8.0) with 80 µL of Protease Inhibitor Cocktail (PIC, Sigma-Aldrich, Merck) and sonicated. After centrifugation at 23428 x g at 4°C for 15 min, the supernatant was collected and incubated with the nickel gel (500 µL) for 2 h at 4°C with vertical rotation. The gel was washed 2x with 10 mL washing buffer (50 mM NaH_2_PO_4_xH_2_O, 300 mM NaCl, pH 8.0) and 2x with washing buffer containing 10 mM imidazole. Before elution the gel was transferred to a 2 mL Pierce™ Centrifuge Columns (Thermo Fischer Scientific) and three elution steps (dropwise) performed with 500 µL elution buffer (50 mM NaH_2_PO_4_xH_2_O, 300 mM NaCl, 250 mM Imidazole pH 8.0). The purified allergens were dialyzed in 2x PBS and stored at 4°C. Bac-Cul o 1P was purified under denaturing condition using the same affinity gel and a comparable procedure as for the native purification with the following changes: The pellet was resuspended in lysis buffer (6 M guanidinium-HCl, 100 mM NaH_2_PO_4_xH_2_O, pH 8.0), the supernatant was incubated for 1 h at RT with the affinity gel and then washed 2x with a wash buffer containing urea (8 M urea, 100 mM NaH_2_PO_4_xH_2_O 10 mM Tris, 100 mM NH_4_Cl, pH 8.0). The bound proteins were eluted in a 2 mL Pierce™ Centrifuge Columns (Thermo Fischer Scientific) with elution buffer (20 mM Tris, 500 mM NaCl, 400 mM L-arginine-HCl, 5 mM β-cyclodextrin, 100 mM Glycerol, 350 mM imidazole), supporting refolding og the protein and dialysed in 2xPBS.

The purified and dialyzed proteins were analysed with protein staining, 2 µg protein/well was loaded on the gel and the gel stained with GelCode^TM^ Blue Safe Protein stain (Thermo Fischer Scientific). Western blot was carried out for confirmation of the proteins, 0.5 µg/well was loaded to the gel and blotted to a PVDF membrane before protein-specific antibodies were applied. The polyclonal antibodies were used at a dilution of 1:5000 except for anti-Cul o 8 (1:2000) and anti-Cul o 11 (1:20000) (supplementary table 1, (31)). For Cul o 7, polyclonal antibody specific for Cul n 4 (14) was used as the two proteins have 33% a.a. identity (15) and the pAb is cross-reactive to Cul o 7. Cul o 3 is an Antigen 5-like protein and is homologous to Cul n 1 (15, 30), their a.a. identity is 70%. The mAb generated in section 2.2 against Cul n1 is cross-reactive to Cul o 3 and was used at dilution 1:25000. For Bac-Cul o 8 mouse anti-His (Biorad) diluted 1:1000 was used as well as mouse anti-Bac Cul o 8.

### 2.4 Horses and sampling

A total of 52 adult horses (5-22 years, 26 males and 26 females) were included in the study. The majority of the horses belonged to the Icelandic breed (n=41) and 27 of them were born in Iceland and imported to continental Europe. The remaining eleven horses belonged to various other breeds (7 Warmbloods, 2 Franches-Montagnes, 1 Quarter Horse, 1 Tinker). All horses were living in Switzerland in areas infested with *Culicoides*, i.e. where IBH was known to occur. Twenty-eight horses (of which 24 were imported from Iceland; suppl. table 2) were affected with IBH. Twenty-four horses (16 of the Icelandic breed, among which three were imported from Iceland) had no history or clinical signs of this condition and served as healthy controls (H). IBH was diagnosed based on the typical clinical signs and a history of seasonal recurring pruritic dermatitis. Almost all IBH horses were treated against IBH either by wearing IBH blankets or use of various topical repellents or creams. No horse included in the study had been treated with systemic corticosteroids in the months preceding blood sampling.

Blood samples were taken by jugular venipuncture using ACD-B and Serum Clot Activator-containing vacutainers (Vacuette®; Greiner, St.Gallen, Switzerland). ACD-B blood was used in the CAST within 12 h of blood sampling. Serum was separated and stored at −80°C until used for testing of allergen specific IgE.

The study was approved by the Animal Experimental Committee of the Canton of Berne, Switzerland (No. BE 14/20).

### 2.5 Cellular Antigen Stimulation Test

The Cellular Antigen Stimulation Test (CAST, Bühlmann Laboratories AG, Allschwil, Switzerland) was performed as described previously (11, 32) with few modifications: After sedimentation of the erythrocytes, the leukocyte rich plasma was collected into a 50 mL propylene tubes and centrifuged at 128 x g for 10 min at RT. The plasma was discarded, and the cell pellet resuspended in stimulation buffer (Bühlmann Laboratories AG) containing 10 ng/mL recombinant equine IL-3 (ImmunoTools, Friesoythe, Germany). Cells were incubated for 40 min at 37°C in 96-well tissue culture plates (Greiner Bio-One GmbH, Kremsmunster, Austria) with anti-equine IgE 134 (0.7 μg/ml; (33)) as positive control, with stimulation buffer only to determine the spontaneous sulfidoleukotriene (sLT) release, with *C. obsoletus* and *C. nubeculosus* whole body extracts (2 μg/ml; (17)) and with the eight insect cell expressed *Culicoides* allergens described above each used at a final concentration of 500 ng/ml. A pool of Bac-Cul o 1P, -Cul o 2P and -Cul o 8 allergens was used for titration on cells from few horses using decreasing allergen concentrations (10, 2, 0.5, 0.1, 0.02 μg/mL). Following incubation, plates were centrifuged at 1000 x g for 4 min at 4°C and the supernatants transferred into a new 96-well microtitre tissue culture plate. The plates were kept at −20°C until assayed within the following one to two weeks. Released sLT was measured using the CAST-ELISA following the manufacturer’s instructions (Bühlmann laboratories AG). For further analysis the spontaneous sLT release was subtracted from the values obtained after stimulation with the allergens.

### 2.6 Serum IgE determination by ELISA

The ELISA was performed using 384 well extra high binding polystyrene microtiter plates (Thermo Fisher Scientific). Fifty μL per well were used in all steps except for the blocking step, where 80 μL were used. Plates were coated (37°C, 2h) with the *Culicoides* r-allergens diluted to 1 μg/mL in 0.2 M carbonate–bicarbonate buffer, pH 9.4 (Thermo Fisher Scientific). After 2 washes with 0.9% NaCl, 0.05% Tween-20, blocking buffer consisting of 5% dried milk powder and 5%Tween® 20 in PBS (pH 7.4, Calbiochem, San Diego, USA) was added and the plates incubated at 37°C for 1 h. All samples were tested for one allergen on the same plate and three different allergens could be tested simultaneously on the same plate. The test sera as well as positive and negative control sera were diluted 1:10 in blocking buffer, added in duplicates to the plates and incubated at 4°C overnight on a shaker. The next day plates were washed 4x and monoclonal anti-horse IgE clone 3H10 (1 μg/mL, (34)) was added to the plates and incubated for 2 h at RT on a shaker. After washing, alkaline-phosphatase-conjugated goat-anti mouse IgG with minimal cross-reactivity to horse serum (Jackson ImmunoResearch Laboratories, Inc, West Grove, USA) diluted 1:2000 in blocking buffer was added to each well and incubated for 1½ h at RT on a shaker. After a final wash, plates were developed with 1.5 mg/mL phosphatase substrate (Sigma-Aldrich, Merck) in 10%diethanolamine (Sigma-Aldrich, Merck), pH 9.8. After 30 min, absorbance was measured at 405 nm using a BioTek synergy H 1 microplate reader (Agilent Technologies; www.agilent.com). Blank corrected OD405 values were used for further analyses of serum IgE (sIgE) levels.

### 2.7 Statistical analyses

As the data did not pass Shapiro-Wilk normality test, non-parametric tests were used. GraphPad Prism 10 was used to plot the graphs of released sLT values and sIgE data for each allergen tested. For comparison of sLT or respectively sIgE values between H and IBH horses, the Mann-Whitney U test was performed in GraphPad Prism. The Wilcoxon test was done for the comparison of the sLT release induced by stimulation of PBL with *C. nubeculosus* and *C. obsoletus* whole body extract.

NCSS 11 was used to perform the following statistical analyses:

Receiver operator characteristic (ROC) curves, with the accuracy of the test represented by the area under the curve (AUC) were used to investigate the capacity of the single allergens to discriminate IBH-affected from H horses in CAST or sIgE.

Spearman rank correlation was used for the comparison of released sLTs (in pg/mL) with sIgE levels (OD405) for each r-allergen.

ROC curves were used to select the optimal threshold values. As cut-off, values giving at least a specificity of 95% at the highest accuracy possible were selected. For each allergen, specific sIgE or sLT values (CAST) were then transformed in positive and negative (above and below threshold level). The 2-sided Fisher’s exact test was used to compare the proportion of IBH-affected and control horses with positive allergen-specific CAST or sIgE results, respectively. To assess the agreement between CAST and sIgE results a kappa test for inter-rater agreement was used. Values <0.40 indicate low association; values between 0.40 and 0.75 medium association; and values >0.75 indicate high association between the two raters. p ≤ 0.05 were considered significant throughout the study.

## 3 Results

### 3.1 Comparison of *Culicoides nubeculosus* extract to *Culicoides obsoletus* extract in cellular antigen stimulation test (CAST)

All horses but two released sLT with the anti-IgE used as positive control, these two non-responders (21) were excluded from the study. There were no significant differences in sLT release with anti-IgE between healthy and IBH horses.

To investigate whether the use of *Culicoides* species present in the environment of the horses, i.e. *C. obsoletus* group (CO) midges, would result in a higher sLT production than laboratory-bred *Culicoides nubeculosus* (CN), PBL from the same 37 horses (22 IBH-affected and 15 health controls) were stimulated with CN and CO extracts. IBH-affected horses produced significantly more sLTs than the healthy control horses with both CN and CO extracts (Figure 1A) and with both extracts all H horses but two, had sLT values below the previously defined threshold of 340 pg/mL (11). Remarkably, the same two healthy horses gave false positive results with both CN and CO. Comparison of the values obtained with these two *Culicoides* extracts in the IBH-affected horses showed a significantly higher sLT production following stimulation with CO (median =1105, range 427-3678 pg/mL sLT) as compared to CN extract (median =925, range 184-3317 pg/mL sLT) in a paired Wilcoxon test (Figure 1B). This resulted in a higher sensitivity of the assay with CO (100% with CO vs 91% with CN) with the same specificity of 87% for both extracts.

**Figure 1:**
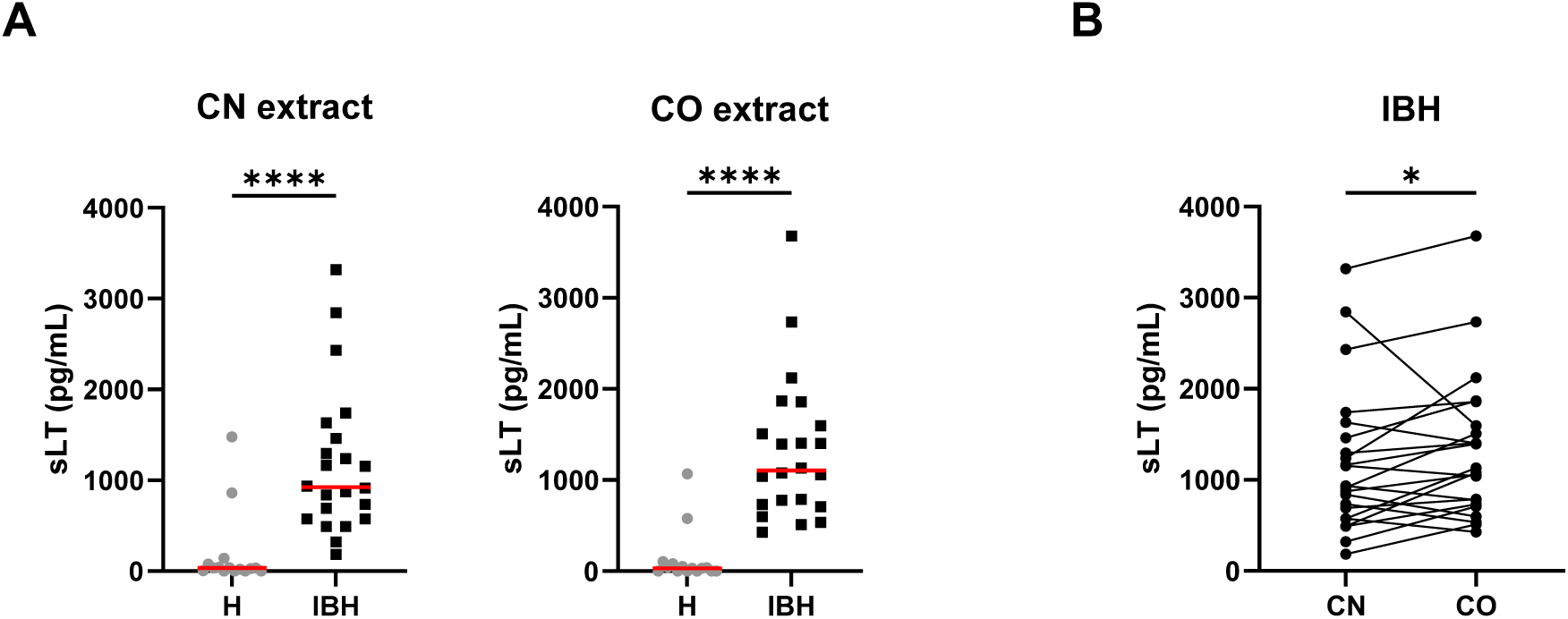
Released sulphidoleukotriene (sLT) following stimulation of peripheral blood leukocytes with *C. obsoletus* (CO) and *C. nubeculosus* (CN) whole body extract. **A)** Concentration of sLT in healthy (H) and IBH-affected (IBH) horses shown as median 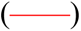 for each group. Dots represent the value of horses. Mann-Whitney U test was used to compare the difference between the groups, **** p ≤ 0.0001. **B)** Concentration of sLT in the IBH-affected horses following stimulation with CN and CO extracts. Dots connected with a line represent the values from a single horse. Comparison between extracts was performed with Wilcoxon test, * p ≤ 0.05.

### 3.2 Expression of *Culicoides* allergens in insect cells

Eight major *Culicoides obsoletus* allergens were cloned into different expression vectors (Table 1) and successfully produced in High five insect cells with the Bac-to Bac Baculovirus expression system (Bac-allergens). All except Cul o 1P were purified under native conditions before dialysed into 2xPBS (Figure 2A). The yield of pure dialysed protein ranges from 1.6 to 4.8 mg from 600×10^6^ High five insect cells, depending on the protein. Bac-Cul o 9 gave the highest yield and Bac-Cul o 7 the lowest.

**Figure 2:**
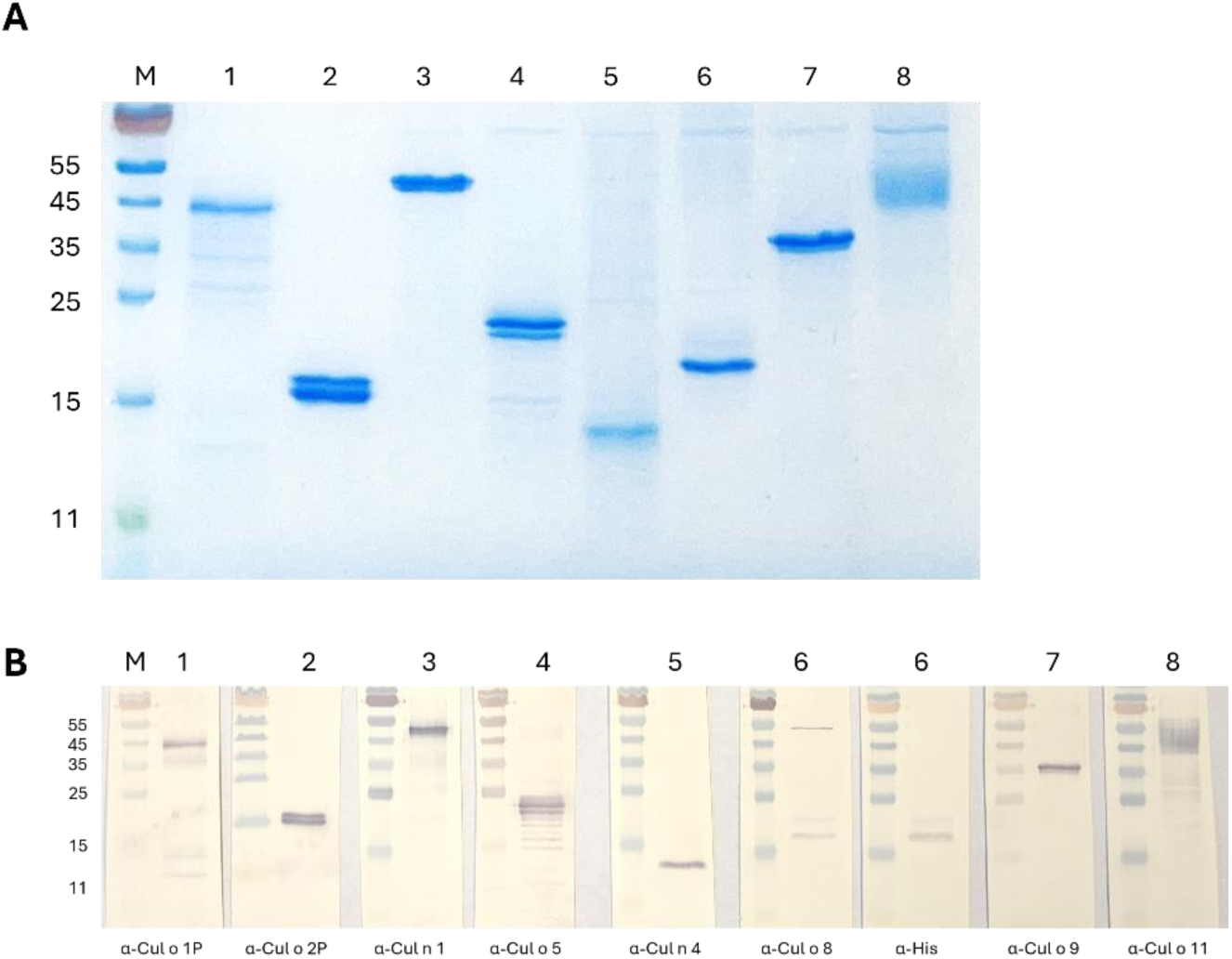
SDS-PAGE of purified and dialysed *Culicoides* allergens expressed in High five insect cells. **A)** Protein staining: 2 µg of each allergen was loaded on the gel. **B) Immunoblot:** 0.5 µg of each allergen was loaded on the gel. **M**: PageRuler marker, **1**: Bac-Cul o 1P, **2**: Bac-Cul o 2P, **3**: Bac-Cul o 3, **4**: Bac-Cul o 5, **5**: Bac-Cul o 7, **6**: Bac-Cul o 8, **7**: Bac-Cul o 9, **8**: Bac-Cul o 11.

Bac-Cul o 2P, -Cul o 5, -Cul o 8 and -Cul o 11 showed protein bands in SDS-PAGE corresponding to the calculated size, while Bac-Cul o 7 appeared as a smaller protein than expected. Bac-Cul o 1P, -Cul o 3 and Cul o 9 appeared as bands approximately 8-10 kDa larger than calculated (Figure 2A).

On the immunoblot the antibodies generated bound to the corresponding proteins seen on the SDS-PAGE (Figure 2 A and B). Anti-Cul o 1P bound to a band close to the 45 kDa marker, anti-Cul o 2P to the double band seen on the SDS-PAGE, at the 15 kDa, anti-Cul o 5 to the double band seen below the 25 kDa, anti-Cul o 8 to the double band above the 15 kDa, the same is seen with the anti-His antibody. Anti-Cul o 9 bound to the single band above the 35 kDa and anti-Cul o 11 to the broad fussy band at 45 to 55 kDa. The polyclonal antibody against Cul n 4 bound to the single band below the 15 kDa when tested on Cul o 7. The monoclonal antibody generated against the Cul n 1 bound to the single band seen between the 45 and 55 kDa when tested on Cul o 3 (Figure 2B).

### 3.3 Titration of Bac-allergens in Cellular Antigen Stimulation Test

PBL from IBH-affected horses were stimulated with a pool of three Bac-allergens Bac-Cul o 1P, -Cul o 2P and -Cul o 8, to determine the optimal concentration of the Bac-allergens to be used in CAST. The median sLT produced following stimulation with 10 μg/mL was 396 pg/mL sLT (Figure 3). It gradually increased with decreasing allergen concentrations and peaked at 0.5 μg/mL (median = 682 pg/mL), before decreasing again at allergen concentrations of 0.1 (485 pg/mL sLT) and even further at 0.02 μg/mL. Depending on the horse, the highest values were obtained with allergen concentrations of 2, 0.5 or 0.1 μg/ml, but the differences between these three allergen concentrations were minimal for the single horses. Punctual testing of these concentrations for the single allergens in few horses confirmed that 0.5 μg/mL was usually the concentration leading to the highest sLT production (data not shown) and this concentration was then used in the subsequent experiments.

**Figure 3:**
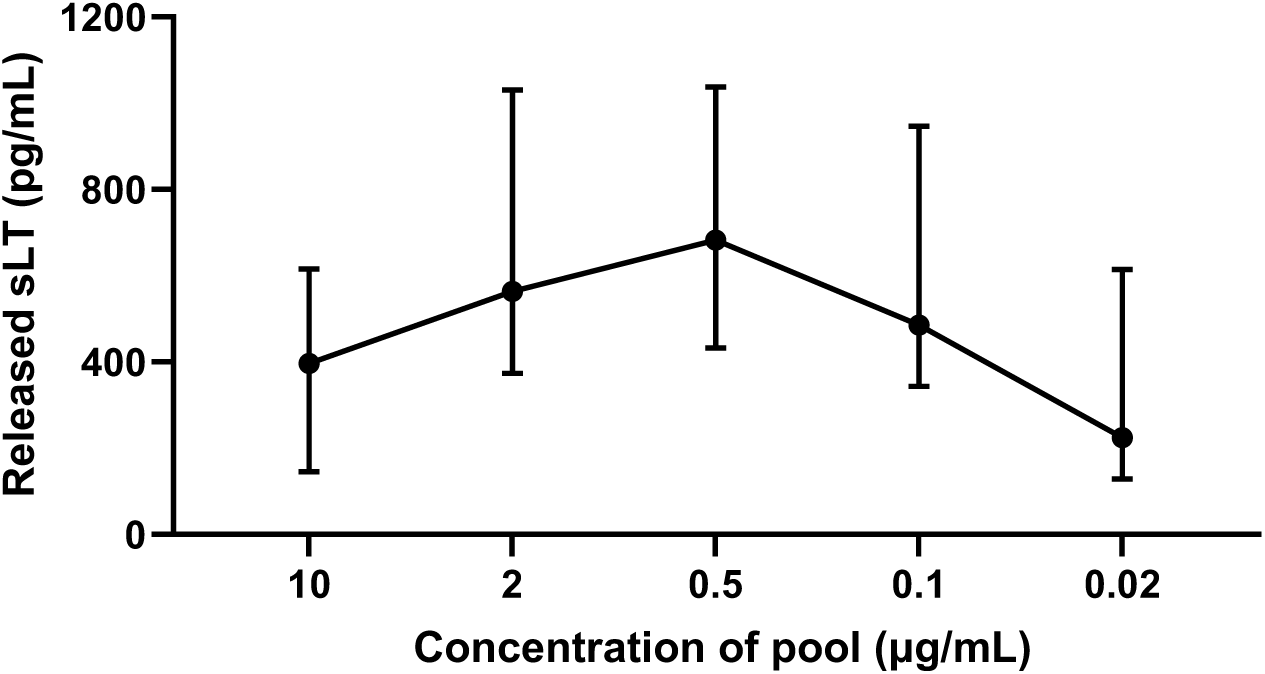
Titration of a pool of insect-cell expressed (Bac-) *C. obsoletus* allergens in cellular antigen stimulation test. Released sulfidoleukotriene (sLT) following stimulation of peripheral blood leukocytes with different concentration of the pool of Bac-Cul o 1P, Cul o 2P and Cul o 8. The sLT is shown for each concentration as median with interquartile range (IQR).

### 3.3 Cellular Antigen Stimulation Test with the single Bac-*Culicoides* allergens

PBL from the 52 horses were stimulated with the single Bac-allergens at the concentration of 0.5 µg/mL and released sLT measured. For all eight Bac-allergens significantly more sLT was released from PBL of IBH-affected horses as compared to healthy horses (Figure 4). The highest median sLT production in the IBH-affected horses was observed after Bac-Cul o 8 stimulation followed by Bac-Cul o 3, -Cul o 9, -Cul o 5, -Cul o 11, -Cul o 7, -Cul o 2P and -Cul o 1P (Figure 4). The sLT release in the H horses was usually < 70 pg/mL.

**Figure 4:**
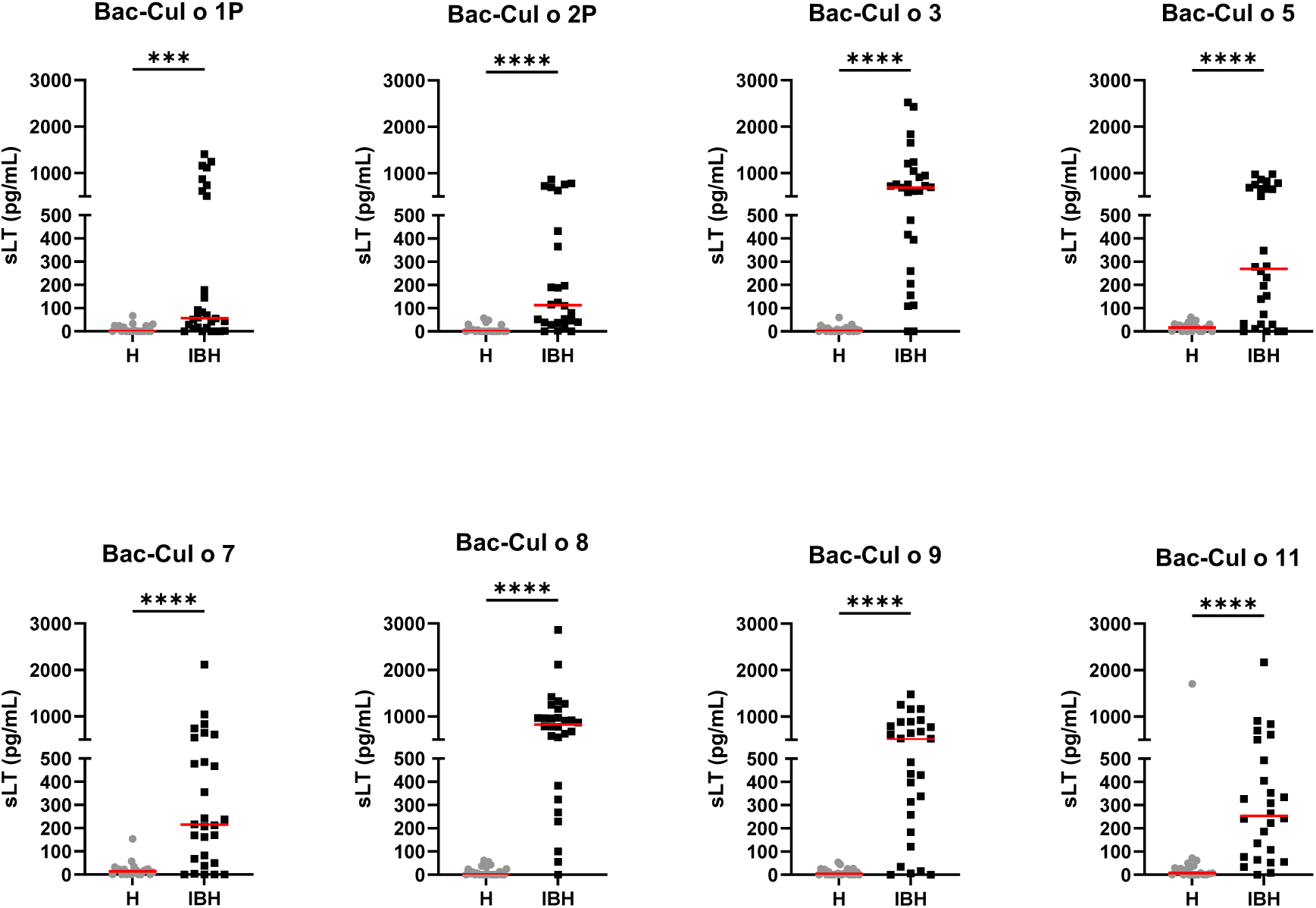
Released sulphidoleukotriene (sLT) following stimulation of peripheral blood leukocytes with the eight insect-cell expressed (Bac-) *C. obsoletus* allergens. Concentration of sLT plotted for healthy (H) and IBH-affected (IBH) horses shown with median for each group in red. Each dot represents the value of single horse. Mann-Whitney U test was used to compare the difference between the groups, *** p ≤ 0.001, **** p ≤ 0.0001.

A ROC curve analysis (Table 2) showed that three of these Bac allergens, namely Cul o 8, Cul o 3 and Cul o 9, had an excellent performance to discriminate IBH-affected from H control horses (AUC > 0.91). The highest AUC was obtained for Cul o 8 (AUC= 0.971, p<0.0001), followed by Bac Cul o 3 (AUC=0.946) and Bac Cul o 9 (AUC=0.919). The performance of the other five Bac allergen was somewhat lower but still good with AUC ranging from 0.809 to 0.892 (p<0.0001). A cut off analysis revealed that for each of the single Bac Cul o allergens a threshold of 100 pg/mL resulted in at least 95% specificity (Table 2) with sensitivities ranging from 36% (Cul o 1P) to 93% (Cul o 8 and Cul o 3) depending on the Bac allergen.

**Table 2:** Overview of the performance of the Cellular Antigen Stimulation Test (CAST) with the eight *Culicoides obsoletus* allergens expressed in insect cells using peripheral blood leukocytes from 28 IBH-affected and 24 control horses. Results from ROC analyses including pairwise accuracy (AUC; upper 1-sided P-value <0.001 for all allergens) with lower and upper confidence limits, and threshold values selected to obtain a specificity ≥ 95% and the resulting sensitivities (Sens) and specificities (Spec).

| Allergen | AUC | Z-Value | 95% confidence limit |  | cut off<br>(pg/ml SLT) | Sens | Spec |
| --- | --- | --- | --- | --- | --- | --- | --- |
|  |  | to test<br>AUC >0.5 | Lower | Upper |  |  |  |
| Bac-Cul o 1P | 0.809 | 5.15 | 0.6549 | 0.8981 | 100 | 0.36 | 1 |
| Bac-Cul o 2P | 0.891 | 8.39 | 0.7545 | 0.9536 | 100 | 0.54 | 1 |
| Bac-Cul o 3 | 0.946 | 11.95 | 0.7984 | 0.9866 | 100 | 0.93 | 1 |
| Bac-Cul o 5 | 0.832 | 5.39 | 0.6652 | 0.9193 | 100 | 0.68 | 1 |
| Bac-Cul o 7 | 0.847 | 5.75 | 0.6787 | 0.9307 | 100 | 0.68 | 0.95 |
| Bac-Cul o 8 | 0.971 | 17.48 | 0.8293 | 0.9954 | 100 | 0.93 | 1 |
| Bac-Cul o 9 | 0.919 | 10.06 | 0.7846 | 0.9708 | 100 | 0.82 | 1 |
| Bac-Cul o 11 | 0.892 | 7.04 | 0.7149 | 0.9615 | 100 | 0.73 | 0.95 |

For Cul o1P and Cul o 2P decreasing the cut off to 50 pg/mL resulted in an increased sensitivity with a still high specificity of 95%. However, because values <100 pg/mL are at the low end of the standard curve in the assay, we feared that technical variability may affect the results when using a threshold <100 pg/ml. We therefore used a threshold of 100 pg/mL for all Bac allergens (Table 2).

### 3.4 Allergen-specific IgE in sera

Sera from the same horses were tested for allergen-specific IgE on the eight Bac-allergens. The IBH-affected horses had significantly higher IgE values against each of the eight Bac-allergens as compared to healthy horses (Figure 5). Specific IgE levels were usually low (OD405<0.1) or not detectable in the sera of the H control horses, except for Cul o 8 and Cul o 11 where OD405 values up to 0.45 or 0.3, respectively, were observed in numerous healthy control horses. ROC analysis shows that the highest AUC were obtained for Cul o 8, Cul o 2P, Cul o 7 and Cul o 9 (AUC> 0.990) (Table 3). The AUC was also high (>0.900) for Cul o 1P, Cul o 3 and Cul o 5 and was only below 0.900 for Cul o 11. A specificity ≥ 95% was obtained at a cut off value of 0.1 (OD405) for all allergens but Cul o 5 (Spec. 92%), Cul o 8 (Spec. 63%) and Cul o 11 (Spec. 67%). Selecting cut off values of 0.5 and 0.3 for Cul o 8 and Cul o 11, respectively, resulted in specificities of 100 and 96% for these two respective allergens. At these given thresholds, a sensitivity of 96% was obtained for Cul o 8 and Cul o 9, followed by Cul o 3 (86%), Cul o 2P and Cul o 5 (82%), Cul o 7 (78%), Cul o 1P (75%) and Cul o 11 (68%) (Table 3).

**Figure 5:**
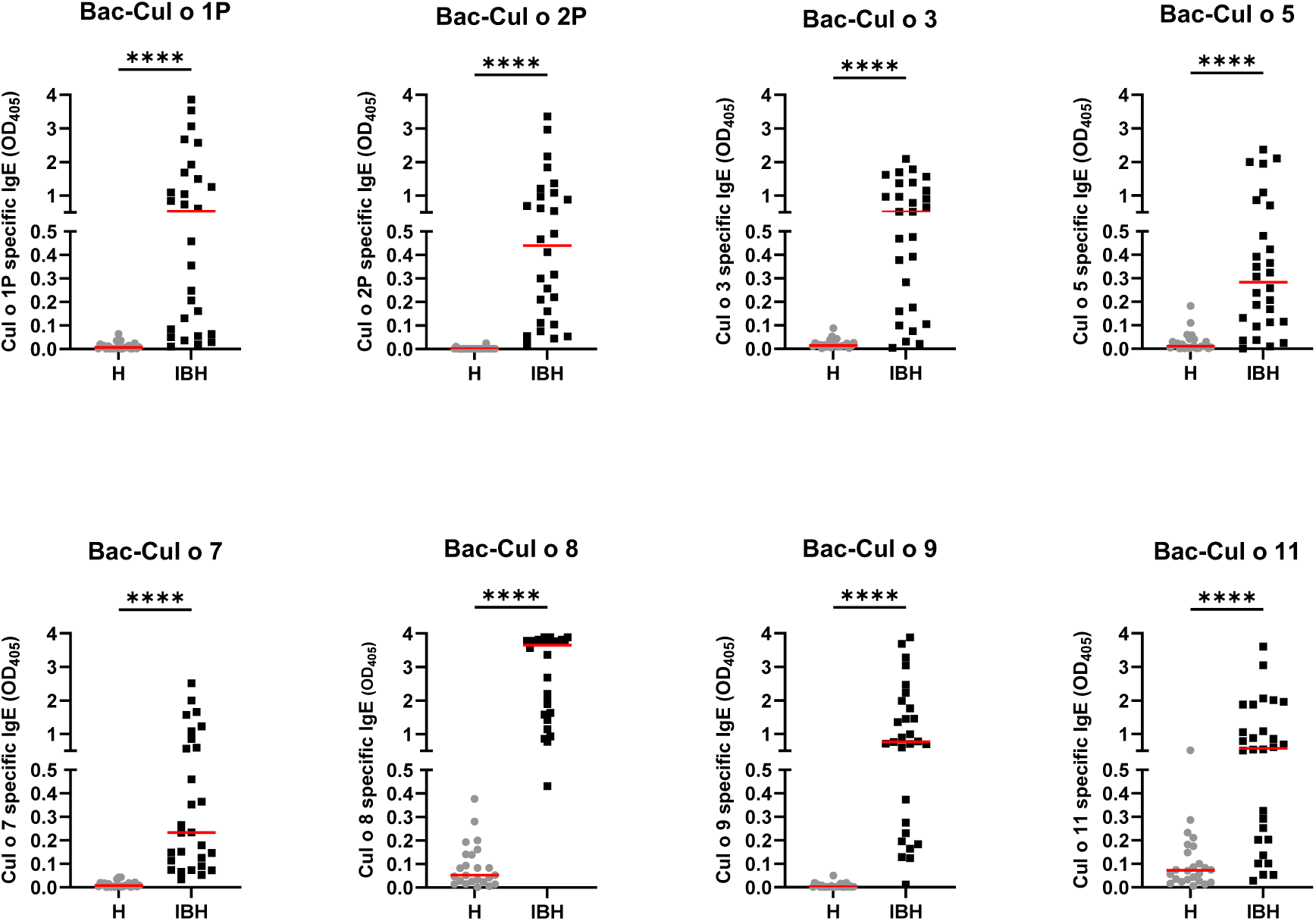
Serum IgE from healthy (H) and IBH-affected (IBH) horses specific for the eight insect-cell expressed (Bac-) *C. obsoletus* allergens tested by ELISA. OD405 value of each horse plotted and shown with the median for the group in red. Mann-Whitney U test was used to compare the difference between the groups, **** p ≤ 0.0001.

**Table 3:** Overview of perfomance of the serum IgE ELISA specific for the eight insect-cell expressed *Culicoides obsoletus* allergens. IgE levels in sera of 28 IBH-affected and 24 control horses. Results from ROC analyses including pairwise accuracy (AUC; upper 1-sided P-value <0.001 for all allergens) with lower and upper confidence limits, and threshold values selected to obtain a specificity ≥ 95% and the resulting sensitivities (Sens) and specificities (Spec).

| Allergen | AUC | Z-Value | 95% confidence limit |  | cut off<br>OD 405 | Sens | Spec |
| --- | --- | --- | --- | --- | --- | --- | --- |
|  |  | to test AUC<br>>0.5 | Lower | Upper |  |  |  |
| Bac-Cul o 1P | 0.968 | 23.89 | 0.896 | 0.990 | 0.1 | 0.75 | 1 |
| Bac-Cul o 2P | 0.999 | 236.88 | 0.976 | 1.000 | 0.1 | 0.82 | 1 |
| Bac-Cul o 3 | 0.946 | 12.54 | 0.810 | 0.986 | 0.1 | 0.86 | 1 |
| Bac-Cul o 5 | 0.907 | 9.27 | 0.772 | 0.964 | 0.1 | 0.82 | 0.92 |
| Bac-Cul o 7 | 0.996 | 97.19 | 0.959 | 1.000 | 0.1 | 0.78 | 1 |
| Bac-Cul o 8 | 1.000 | 4.5403E+15 |  |  | 0.5 | 0.96 | 1 |
| Bac-Cul o 9 | 0.994 | 75.23 | 0.949 | 0.999 | 0.1 | 0.96 | 1 |
| Bac-Cul o 11 | 0.878 | 7.98 | 0.745 | 0.944 | 0.3 | 0.68 | 0.96 |

### 3.5 Comparison of CAST and IgE ELISA

The CAST and the IgE ELISA were compared using Spearman’s rank correlation and kappa coefficients. Table 4 shows that there were significant (p<0.0001) positive correlations between released sLT in CAST and serum IgE values for each of the eight *Culicoides* allergens, ranging from 0.610 (Cul o 1P) to 0.818 (Cul o 8). Comparison of the categorical CAST and serum IgE results showed an excellent agreement for Cul o 8 and Cul o 9 (kappa ≥0.85) and a substantial agreement for Cul o 7, Cul o 5, Cul o 3 and Cu o 11 (0.61>k<0.8). The agreement was lower for Cul o 2P and Cul o 1P (Table 4).

**Table 4:** Comparison of sLT production measured in CAST and serum IgE levels for the eight insect-cell expressed *Culicoides obsoletus* allergens determined in 52 horses.

| Allergen | Spearman<br>correlation<br>coeff | Kappa | Kappa<br>2-sided<br>prob |
| --- | --- | --- | --- |
| Bac-Cul o 1P | 0.610 | 0.54 | <0.000 |
| Bac-Cul o 2P | 0.768 | 0.48 | <0.00 |
| Bac-Cul o 3 | 0.759 | 0.73 | <0.0000 |
| Bac-Cul o 5 | 0.738 | 0.75 | <0.0000 |
| Bac-Cul o 7 | 0.791 | 0.79 | <0.0000 |
| Bac-Cul o 8 | 0.815 | 0.96 | <0.0000 |
| Bac-Cul o 9 | 0.818 | 0.85 | <0.0000 |
| Bac-Cul o 11 | 0.74 | 0.68 | <0.0000 |

For further comparison of the two assays, the positivity rate of the groups was plotted for each allergen (Figure 6). For all allergens, except Cul o 1P and 2P, the positivity rates were similar in both assays, with usually a slightly higher positivity in IgE serology compared to CAST. Bac-Cul o 8, -Cul o 3 and -Cul o 9 showed the highest positivity rates in the IBH group in both assays, followed by Bac-Cul o 5, -Cul o 7 and Cul o 11. High differences in positivity rates in the assays were observed for Bac-Cul o 1P and Cul o 2P, with a markedly higher positivity in the IgE ELISA (71.4% and 82.1%, respectively) compared to the CAST (35.7% and 53.9%, respectively). Decreasing the cut off to 50 pg/mL sLT increased the sensitivity in the CAST to 57% and 65%, for Cul o 1P and Cul o 2P, respectively. Healthy horses were negative in both assays, except in CAST on Bac-Cul o 7 and -Cul o 11, and in IgE ELISA on Bac-Cul o 5 and -Cul o 11, resulting in a small proportion of false positive results (Figure 6). The overall data from the healthy horses showed a very good agreement between CAST and IgE test results (Figure 7A), as 98% of the healthy horses were negative in both. Few discordant results were obtained for Cul o 5, Cul o 7 and Cul o 11. The overall data from the IBH-affected horses was more complex (Figure 7B). Sixty five percent were positive and 13% negative in both tests, resulting in an overall agreement of 78%. Sixteen per cent of the IBH horses were only positive in IgE serology and 6% were only positive in CAST. For Cul o 8 the agreement was excellent (κ=0.96): all IBH horses were positive in both tests except one, that was only positive in IgE serology.

**Figure 6:**
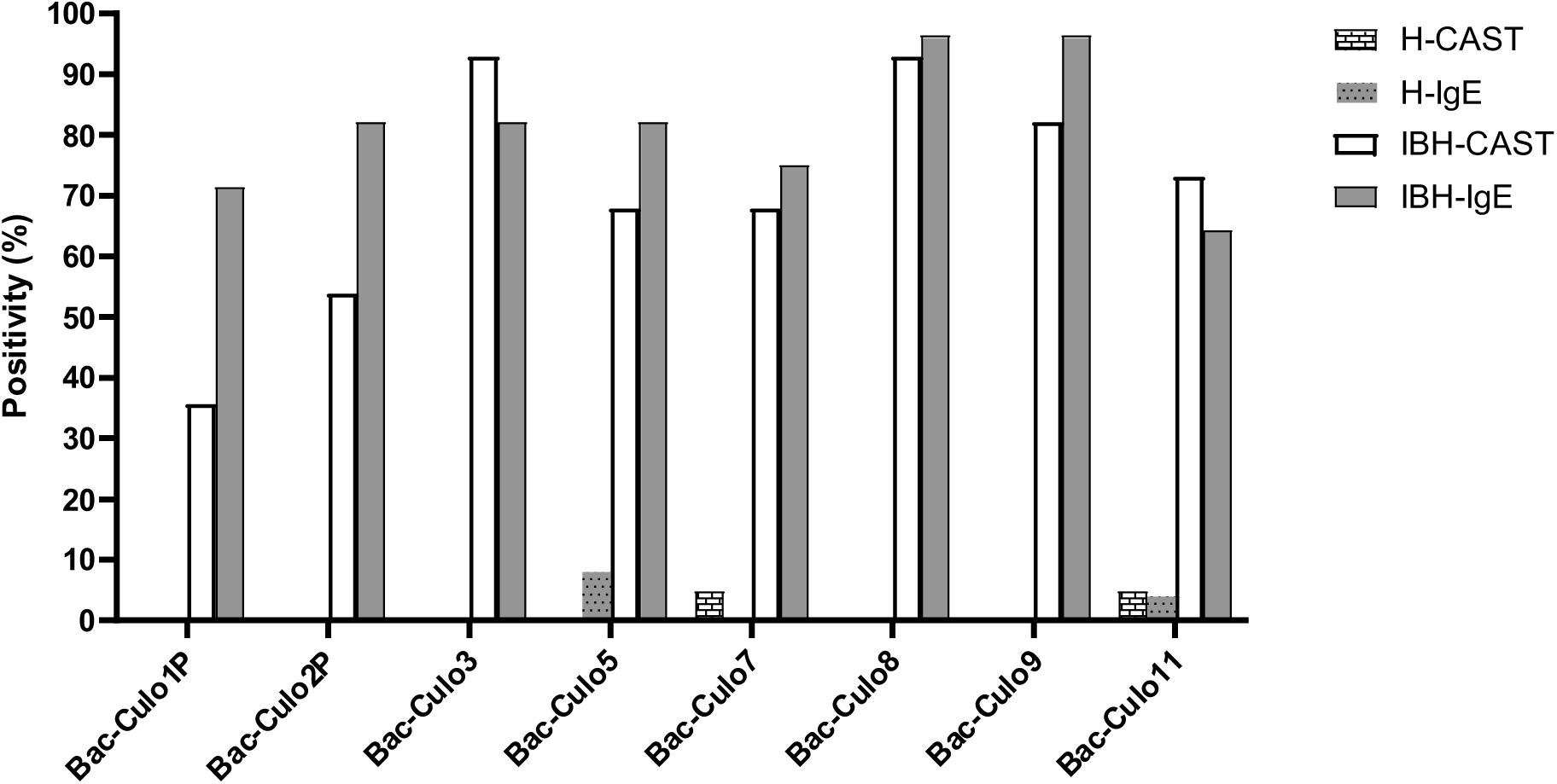
Percentage of horses positive in CAST and IgE ELISA on the eight insect-cell expressed *Culicoides obsoletus* allergens. Percentage of horses positive in CAST (white columns) and in IgE ELISA (grey). Data from healthy horses shown as columns with patterns.

**Figure 7:**
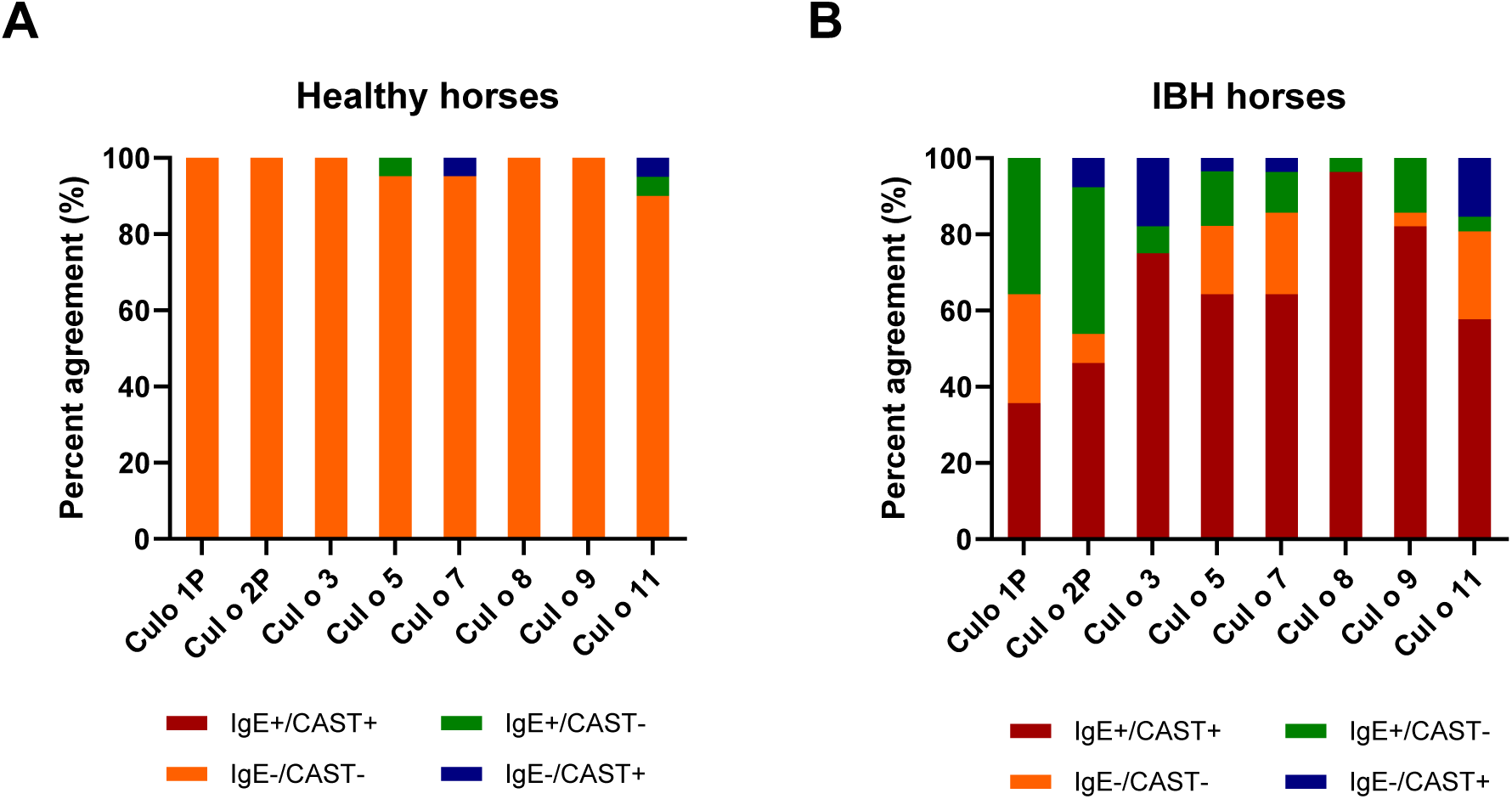
Agreement between CAST and IgE positivity. Agreement between CAST and IgE positivity for the eight *Culicoides obsoletus* allergens. **A)** H control horses. **B)** IBH-affected horses. The red and orange colors show full agreement (CAST+/IgE+ and CAST-/IgE-, respectively). Green: CAST-/IgE+ and blue: CAST+/IgE-.

IBH-affected horses were positive on a median number of 6 recombinant allergens in the CAST and 7.5 allergens in IgE serology. In IgE serology, 42.9% of the IBH horses were positive on the eight tested allergens, followed by 21% on seven allergens. Between 3.6 and 10.8% were positive on two to six allergens (Figure 8A). All IBH horses were IgE positive to at least two allergens. In the CAST, all IBH horses were positive with at least one allergen (Figure 8B). The majority (29.2%) was sensitized to seven allergens and 17% were positive on all eight allergens. Between 4.2 (one horse) and 16.7% were positive with one to six allergens, with only one horse positive to a single allergen. In contrast, all but one (95.2%) H horse were negative in the CAST and this horse was positive with two allergens. In IgE serology 91.7 % of the H horses demonstrated negative test results with the eight allergens, one horses gave a positive result with one allergen and one horse with two allergens (Figure 8A).

**Figure 8:**
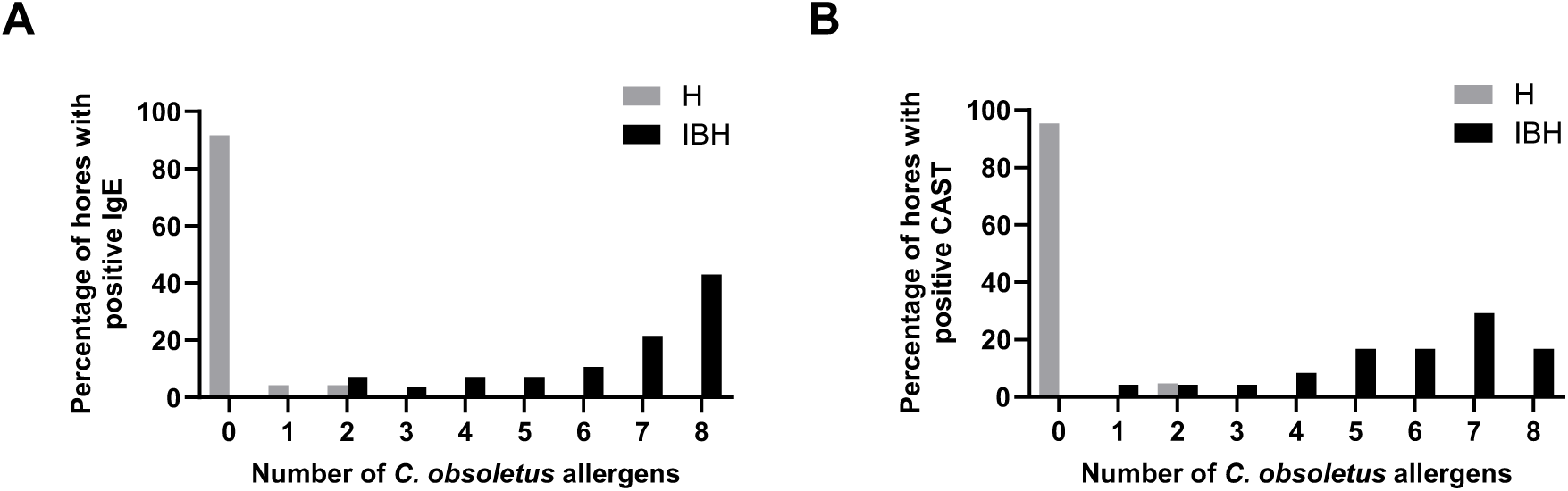
Cumulative positivity to the eight *C. obsoletus* recombinant allergens in IgE serology (A) and CAST (B). Percentages of IBH-affected and healthy horses against the number of recombinant allergens to which they react.

## 4 Discussion

Insect bite hypersensitivity is the most common and best characterized IgE-mediated allergy in horses and is considered a natural model of allergy (35). However, the limited availability of horse specific assays and the absence of pure and functional allergens has hindered the development of new and more effective allergen immunotherapy strategies. The identification of the specific *Culicoides* allergens and their production as pure recombinant proteins, replacing crude *Culicoides* whole body extracts, has been a significant advancement for research (reviewed in 36, 37) and for the development of more reliable serological IgE tests for diagnosing IBH. However, most *Culicoides* allergens have been expressed in *E. coli* and the majority could not be purified and refolded under native conditions. This limitation has restricted their usefulness in cellular assays (18) particularly in sulfidoleukotriene release assays. In the present study, expression in insect cells has allowed for the purification of all but one (Bac-Cul o 1P) under native conditions. Nevertheless, all proteins including Bac-Cul o 1P could be dialyzed into 2x PBS suggesting that the Bac-Cul o 1P was probably refolded correctly.

The protein specific antibodies used for confirmation of the Bac-proteins were all generated using *E. coli* expressed proteins, except for Cul o 8, where mice were immunized with Bac-Cul o 8. The protein band seen with the anti-Bac Cul o 8 antibody was confirmed using anti-His antibody. The size of the protein bands was in line with calculated size of the construct for Cul o 2P, Cul o 5, Cul o 8 and Cul o 11. For Cul o 1P, Cul o 3 and Cul o 9 the protein band seen with protein staining and in immunoblot were larger than the calculated size. The genes encoding for those three proteins were cloned into pI-secSUMOStar vector and therefore have a SUMO-tag. The calculated size of the SUMO-tag is 11 kDa but it appears as a 18 kDa band after cleavage. With this addition, the protein bands correspond to the expected protein size.

Bac-Cul o 7 appears as a smaller band than expected. The *Cul o 7* gene was cloned into FastBac1 with His-tag at the C-terminus. Cul o 7 protein has a predicted secretion signal (a.a. 1-24) which could have been cleaved, resulting in a smaller mature protein, as detected in the SDS-PAGE and immunoblot.

Of the numerous *Culicoides* allergens described so far (reviewed in 37) the selection of allergens for expression in insect cells was based on the demonstrated higher relevance of *C. obsoletus* over *C. nubeculosus* allergens as shown in the CAST (Figure 1) and confirming previous studies (12). Eight of the nine previously identified major *C. obsoletus* allergens were expressed in insect cells. Unfortunately, the nineth allergen, Cul o 10, could not be produced in time due to technical challenges. This study demonstrates that the eight major *C. obsoletus* allergens expressed in insect cells are functional allergens for IBH as they induce sLT release in IBH-affected horses. Notably, very low sLT levels were detected in samples from healthy horses, enabling the use of a low threshold of 100 pg/mL, which is below the value (200 pg/mL) usually recommended by the manufacturer. While the CAST and IgE serology measure different stages of the allergic cascade, there was excellent agreement between sLT release and sIgE levels for most allergens, except for Cul o1P and Cul o 2P. For Cul o 1P fewer IBH horses were CAST-positive compared to IgE serology. This discrepancy may stem from the fact that Cul o 1P could not be purified under native conditions, potentially impairing its ability to crosslink cell-bound IgE molecules. Lowering the CAST threshold did not substantially improve agreement with IgE serology (κ=0.61) for this allergen as also indicated by the moderate correlation between sLT and sIgE. In contrast, for Cul o 2P the correlation between sLT release and sIgE values was higher (0.77) but agreement between tests was low (κ=0.48) when using the 100 pg/ml threshold in the CAST. Reducing the threshold to 50 pg/ml sLT improved slightly the agreement between both tests (κ=0.57) as well as CAST sensitivity (65%). However, to account for potential technical variability we found it more reliable to use a cut off that was not below 100 pg/ml. For diagnostic purposes, using the three best-performing allergens in CAST, Cul o 8, Cul o 3 and Cul o 9 could be sufficient: Combining the results of these three allergens yielded 100% sensitivity and specificity, suggesting that they could replace *C. obsoletus* whole body extract in the CAST. Production of *C. obsoletus* extract is very cumbersome as the midges have to be captured from the wild, while the use of laboratory-bred *C. nubeculosus* extract results in a lower performance of the CAST (Figure 1). Thus, the use of few recombinant *C. obsoletus* allergens represents a substantial improvement of this assay for diagnosing IBH. Inclusion of the eight *Culicoides* recombinant allergens in the CAST should be considered for allergen selection for potential molecular-based, patient tailored immunotherapy.

The sIgE ELISA using these insect cell expressed allergens exceeded expectations, showing very high sensitivity and good specificity (87%) when combining the results from all allergens. Sensitization to allergens without clinical signs is well-known to occur (38). Accordingly, a relatively high number of H horses exhibited sIgE against Cul o 8 and/or Cul o 11. This may be explained by true sensitization or by cross-reactive carbohydrate determinants (CCDs) especially for Cul o 11 which is strongly glycosylated as evidenced by the broad band in the SDS-PAGE and immunoblot (Figure 2A and B) that resolved to a lower molecular weight band after deglycosylation (39). This was also observed albeit to a lesser extend to Cul o 8 (39). Interestingly, high thresholds were also required in Novotny et al. for these allergens, even when expressed in *E. coli,* suggesting true sensitization in some healthy horses (17). Further studies are needed to investigate the effects of glycosylation on sIgE reactivity and whether CCD blockers can reduce IgE-binding in healthy horses. Nonetheless, adjusting the ELISA threshold improved specificity without notable loss of sensitivity. Both assays performed above expectations, emphasizing the importance of using pure and correctly folded allergens. Another contributing factor may be that the IBH group predominantly comprised horses imported from Iceland (suppl. table 2). Icelandic horses born in Iceland and imported to Europe have a high degree of sensitization to *Culicoides* allergens when developing IBH compared to IBH horses born in continental Europe affected with IBH (17). Unfortunately, the number of IBH horses in this study was too small to evaluate differences between horses born in Iceland and horse born in continental Europe. Importantly, while functional allergens such as those described here are invaluable for research, they are probably unsuitable for AIT, due to the risk of mast cell degranulation and the resulting potentially severe side-effects. *E.coli* produced allergens represent a safer alternative for AIT (40).

Our findings confirm the high relevance of these eight allergens for IBH, as previously identified by Novotny et al. using microarray technology (17). While microarrays are efficient for testing many allergens simultaneously with minimal serum and reagents, ELISAs are better suited for testing limited number of allergens. The results presented here indicate that the sIgE ELISA with the eight *Culicoides obsoletus* allergen can be standardized for reliable IBH diagnostics, while the CAST appears more robust for confirming the absence of allergy, for example in pre-purchase examinations or in cases of discordant clinical signs and IgE serology.

In conclusion, insect cell expressed *Culicoides* recombinant allergens provide new opportunities for studying Culicoides hypersensitivity not only in horses, but also potentially in human patients. Functional *in vitro* assays such as the CAST with Bac-Cul o allergens will be valuable for monitoring the response to allergen immunotherapy and for refining immunotherapy protocols in this natural allergy model.

## Author Contribution

SJ: Conceptualization, formal analysis, funding acquisition, project administration, visualization, writing - original draft, review and editing. SBS: Investigation, review and editing. JM: Investigation, validation, review and editing. AZ: Investigation, resources, validation, review and editing. ST: Funding acquisition, methodology, investigation, review and editing. EM: Conceptualization, data curation, formal analysis, funding acquisition, project administration, supervision, writing - original draft, review and editing.

## Acknowledgments

We are thankful to Dr. Bettina Wagner at the Department of Population Medicine and Diagnostic Sciences, College of Veterinary Medicine, Cornell University, Ithaca, NY 14853, USA for providing the anti-IgE antibody, clone 134.

---

**Supplementary table 1:** List of *Culicoides* antigens used for production of specific antibodies in mice.

| Antigen used | Produced | Type of mouse<br>antibody | Dilution used<br>in WB |
| --- | --- | --- | --- |
| *E.coli* -Cul o 1P | Keldur | pAb | 1/5000 |
| *E.coli* -Cul o 2P | Keldur | pAb | 1/5000 |
| *E.coli* -Cul n 1* | Keldur | mAb | 1/25000 |
| *E.coli* -Cul o 5 | ArcticLas | pAb | 1/5000 |
| *E.coli* -Cul n 4** | Keldur | pAb | 1/5000 |
| Bac-Cul o 8 | ArcticLas | pAb | 1/2000 |
| *E.coli* -Cul o 9 | ArcticLas | pAb | 1/5000 |
| *E.coli* -Cul o 11 | ArcticLas | pAb | 1/20000 |

\*Cross-reactive with Cul o 3
\*\*Cross-reactive with Cul o 7

**Supplementary table 2:**
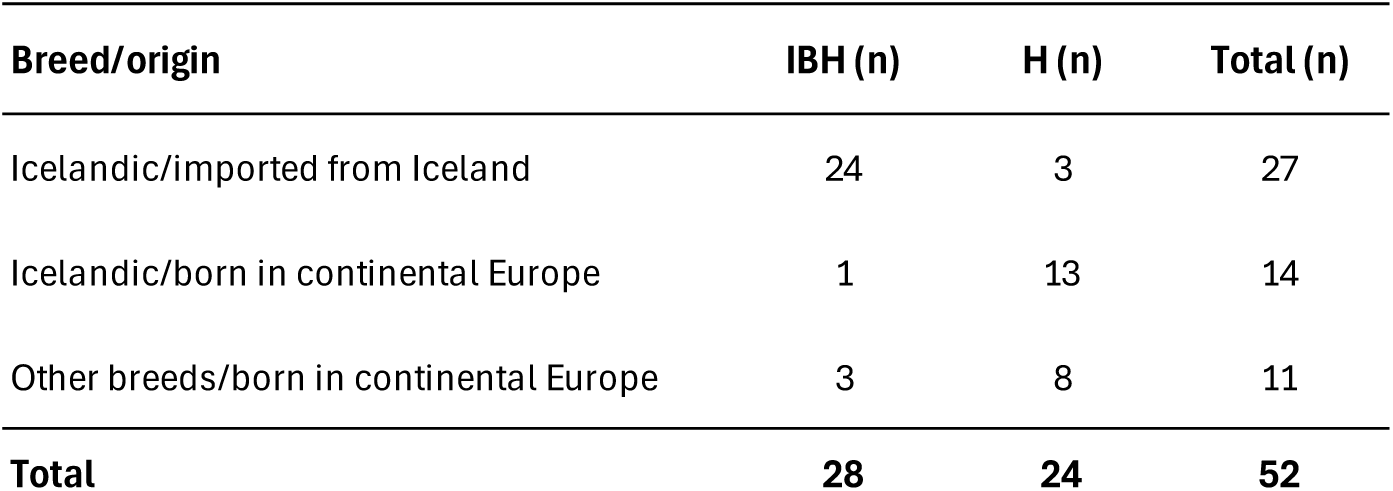
Breed and origin of the horses used in the study.

| Breed/origin | IBH (n) | H (n) | Total (n) |
| --- | --- | --- | --- |
| Icelandic/imported from Iceland | 24 | 3 | 27 |
| Icelandic/born in continental Europe | 1 | 13 | 14 |
| Other breeds/born in continental Europe | 3 | 8 | 11 |
| **Total** | **28** | **24** | **52** |

